# Predicting *in vivo* concentrations of dietary hop phytoestrogens by physiologically based kinetic modeling

**DOI:** 10.1101/2024.08.23.609337

**Authors:** Maja Stevanoska, Karsten Beekmann, Ans Punt, Shana J. Sturla, Georg Aichinger

**Affiliations:** Department of Health Sciences and Technology, ETH Zurich, Switzerland; Wageningen Food Safety Research (WFSR), Netherlands

**Keywords:** endocrine disruption, estrogenicity, polyphenols, kinetic modeling, hops supplements, phytoestrogens

## Abstract

Hop extracts containing prenylated polyphenols such as 8-prenylnaringenin (8-PN) and its precursor isoxanthohumol (iXN) are popular among women seeking natural alternatives to hormone therapy for postmenopausal symptoms. Due to structural similarities with estrogens, these compounds act as estrogen receptor agonists. Especially 8-PN, described as the most potent phytoestrogen known to date, poses a potential risk for endocrine disruption. Therefore, its use as a hormone replacement raises concerns for human health. However, a significant challenge in assessing the potential endocrine-disruptive effects of hop polyphenols is the lack of data on their toxicokinetics. Particularly, information on *in vivo* concentrations in target tissues is lacking. To address this gap, we developed a physiologically based kinetic (PBK) model tailored to female physiology. The model was used to predict the levels of hop polyphenols in human blood and target tissues under realistic exposure scenarios. The predictions suggest that iXN and 8-PN concentrations in target tissues reach the low nanomolar range after dietary supplementation. This study enhances our understanding of the safety profile of hop polyphenols and highlights the need for further research into their use as an alternative to hormone therapy in menopausal women.

## 1. Introduction

Although hormone therapy is a primary recommended treatment for early menopausal symptoms (1), the associated increased risk of breast cancer (2) has led many women to seek natural alternatives (3). Supplements derived from female hop flowers, which are rich in prenylated polyphenols, including chalcones and flavonoids are a popular choice among postmenopausal women to alleviate postmenopausal symptoms (4, 5). However, due to the structural similarity of these phytochemicals to endogenous estrogens, some can bind to estrogen receptors (ERs), initiating direct estrogenic effects and potentially causing endocrine disruption (6). Currently, it remains unclear which physiological levels of these compounds potentially cause endocrine disruption and it is crucial to understand the exposure doses needed to achieve these levels.

The most abundant polyphenol in the hop flower itself is xanthohumol (XN); it is marketed in supplement form for its supposed antioxidant, anti-inflammatory, and *in vitro* anti-tumor properties considered to arise from its activity in modulating tumor-promoting cell signaling (7). However, these hop polyphenols are biochemically interconvertible, and their composition depends on the product preparation process (8). During beer brewing, thermal processing converts XN into isoxanthohumol (iXN), which is a weak estrogen receptor agonist (9). Additionally, the remaining XN is partially converted to iXN in the stomach through acid-catalyzed cyclization (10). While iXN is only a weakly potent estrogen receptor agonist, it can be metabolically activated by hepatic CYP1A2 to the potent phytoestrogen 8-prenylnaringenin (8-PN), in addition to lesser extents of the latter being also present in hops (Figure 1) (11). The amount of 8-PN present in beer varies widely, ranging from 1 µg/L reported in European lager beer and up to 240 µg/L in American porter beer (12). 8-PN is the most potent phytoestrogen known to date, with its affinity to ERs being an estimated 100 times higher than the soy isoflavone genistein and only seventy times lower than the endogenous 17 β-estradiol (13). 8-PN has a more than 2-fold higher affinity for ERα than for ERβ (13). The two receptors have different effects on cell signaling in various tissues, for example, ERα activation is associated with cell proliferation and Erβ activation is associated with counteracting ERα-stimulated cell proliferation (14). ERα is the predominant isoform in the uterus, and it is highly expressed in the endometrium, ovaries, mammary gland, and bones. Therefore, ERα activation by 8-PN may interfere with the regulation of metabolism and endocrine processes (15).

**Figure 1.**
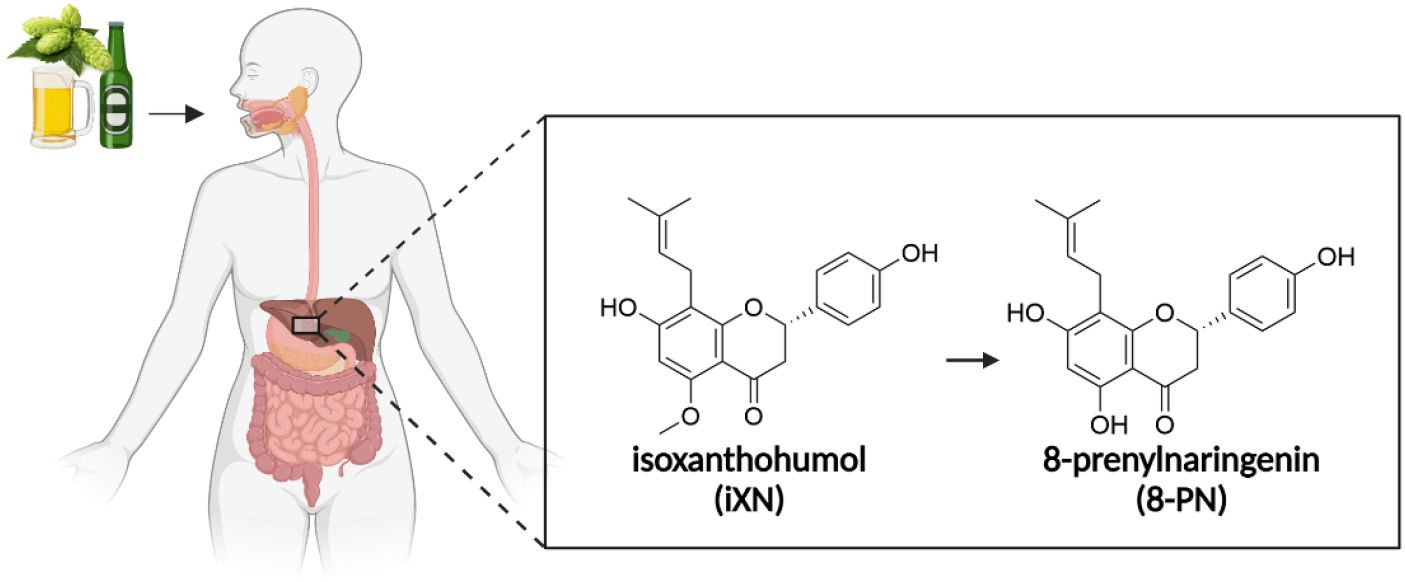
Schematic representation of biotransformation of isoxanthohumol to 8-prenylnaringenin, including their respective chemical structures.

Given the endocrine activity of 8-PN, there is a concern about its potential endocrine-disrupting properties. So far, research has primarily focused on its beneficial effects as an antioxidant and potential natural hormone replacement, with limited studies addressing its potential adverse impacts, as recently reviewed (16). For example, Overk *et al*. evaluated the estrogenic effects of 8-PN in an adult rat model assessing its impact on uterus weight, demonstrating a clear dose-response relationship (17). Specifically, the middle and the highest doses of 8-PN in the study, (4 and 40 mg/kg/d) exhibited significant estrogenic activity, resulting in increased uterine weight. Although the estrogenic activity of 8-PN has been demonstrated, a gap in knowledge remains regarding its effects in human target tissues including when 8-PN is combined with other hop polyphenols and endogenous estrogens.

Accurately predicting *in vivo* outcomes from *in vitro* methods remains a challenge, in part due to the compounds’ complex toxicokinetic parameters (absorption, distribution, metabolism, and excretion; ADME), which play a crucial role in determining target organs of potential toxicity and the time to reach an internal concentration of the compounds of interest, sensitive exposure windows, species, and sex differences (18). There are several available *in vitro* methods to test endocrine activity (19), however, concentrations of chemicals tested *in vitro* often do not correspond with concentrations at the target sites *in vivo*, raising questions about the relevance of the results to realistic exposure scenarios.

Understanding the toxicokinetics of iXN and 8-PN is essential for predicting their systemic and tissue concentrations and estimating the concentration at target sites where toxicity would occur. Therefore, the aim of this study was to enable prediction of systemic and tissue concentrations of iXN and 8-PN upon ingestion of hops supplements containing both iXN and 8-PN by developing a corresponding physiologically based kinetic (PBK) model that uses human physiology, physicochemical properties, and *in vitro* metabolism data. We established a PBK model specifically parameterized to reflect female physiology, which is physiologically relevant for these exposures. The new model facilitates the accurate prediction of blood and tissue concentrations of iXN and 8-PN, particularly in the context of high estrogen receptor expression, providing insight into their safety and potential endocrine activity.

## 2. Methods

### 2.1. Chemicals and reagents

Pooled Human Liver S9 (mixed gender and age donors), isoxanthohumol, 8-prenylnaringenin, uridine-5’-diphosphoglucuronic acid (UDPGA) trisodium salt, magnesium dichloride, *β*-glucuronidases (from bovine liver, 100000 units), and sodium acetate were purchased from Sigma-Aldrich (Buchs, Switzerland). A pooled human intestinal S9 fraction (19 donors, mixed gender) was purchased from Biopredic (Saint-Grégoire, France). Alamethicin was purchased from Enzo Life Sciences AG (Lausen, Switzerland). DMSO was purchased from VWR (Dietikon, Switzerland), and acetonitrile (HPLC-grade) was purchased from Merck-Millipore (Schaffhausen, Switzerland). Dulbecco’s Modified Eagle Medium, fetal bovine serum, HBSS, HEPES, and penicillin/streptomycin were purchased from Gibco, Life Technologies Limited (Paisley, UK).

### 2.2. Glucuronidation kinetics

To quantify hepatic phase II metabolism of iXN and 8-PN, the glucuronidation rates catalyzed by pooled human liver and intestinal S9 fractions of mixed age and gender donors were measured. The incubation mixtures had a final volume of 160 µL and contained 10 mM UDPGA in 50 mM Tris-HCl pH 7.4, 10 mM MgCl_2_, 0.1 mg/mL hepatic or 0.3 mg/mL intestinal S9 fraction in 0.1 M Tris HCl pH 7.4. The incubation mixtures were pre-incubated without the substrate for 5 min at 37 °C with a final concentration of 25 µg/mL alamethicin to increase membrane porosity. To initiate the reaction, the substrates were added at 100 times the concentration in DMSO. All mixtures were incubated for 15 min at 37 °C except the intestinal S9 8-PN mixtures, which were incubated for 5 min. Rate constants for the glucuronidation reaction were determined using final substrate concentrations ranging from 0.25 µM to 100 µM. To terminate the reaction 1:1 ratio of ice-cold methanol was added to the samples. Due to the unavailability of reference glucuronides, additional peaks observed in the HPLC analysis after S9 incubations were confirmed to result from glucuronidation by adding 50 µL of the samples to 0.1 M sodium acetate (pH 5) that contained 1000 units of β-glucuronidase. These mixtures were incubated for an additional 1 h at 37 °C and terminated by the addition of the corresponding volume of ice-cold methanol. Additionally, unterminated samples were incubated with 0.1 M sodium acetate as a control. Samples were kept at -20 °C for 1 h to allow proteins to precipitate. Before analysis, the S9 samples were centrifuged at 20’ 817 g for 5 min at 4 °C. The resulting samples were analyzed by HPLC, as described in section 2.2. Curve fitting of the data to Michaelis-Menten kinetics and the calculation of kinetic parameters were conducted using GraphPad Prism 10 software. The glucuronidation rates were scaled to the human organism based on the mg of S9 protein to g of liver and small intestine with scaling factors commonly used for PBK modeling (VLS9 = 107.3 mg S9 protein/g liver; (20) (VGS9 = 35.2 mg S9 / g small intestine) (21).

### 2.3. HPLC analysis

Original compounds and their corresponding glucuronides were quantified using HPLC-DAD analysis on an Agilent 1200 series instrument. The HPLC system was equipped with a Waters XBridge BEH 130 C18 column, 3.5 μm particle size, 4.6 × 150 mm, and a DAD detector detecting at 295 nm. The liquid phase consisted of water (A) and acetonitrile (B), each supplemented with 0.1% formic acid. The following gradient profile was used at a flow rate of 1 mL/min: 0 min, 20% B; 0-2 min, 50% B; 10-14 min, 80% B; 14-15.5 min, 20% B; post-time 3 min. Reference standards of iXN and 8-PN were used to generate corresponding calibration curves for quantification.

### 2.4. PBK model conceptualization

The PBK model for hop polyphenols developed in this study was based on a previously published model for urolithin A and its glucuronide (22), which was slightly modified and adapted to include organs of interest for estrogenic activity, such as the uterus. The model consists of blood, adipose, liver, gut, uterus, and kidney tissue as separate compartments (Figure 2). The implementation of the uterus as a compartment was based on Teeguarden *et al*. (23), where the uterus partition coefficient was assumed to be the same as the muscle partition coefficient, as the uterus is primarily composed of muscle tissue. Other organs were grouped as slowly perfused tissues (bone, skin, and muscle) and quickly perfused tissues (heart, brain, and lungs). Uptake of iXN was modeled to occur via the small intestinal lumen, and excretion was modeled to occur via the kidneys. As both iXN and 8-PN undergo glucuronidation in the liver and small intestinal tissue, the glucuronidation rates were measured as described above and implemented in the model. A separate submodel was created to predict the levels of iXN and 8-PN glucuronide (iXNGluc, 8-PNGluc) and 8-PN. Additionally, the conversion rate of iXN to 8-PN in the liver was modeled as described below. The main model was extended to three submodels, for iXNGluc, 8-PNGluc, and 8-PN.

**Figure 2.**
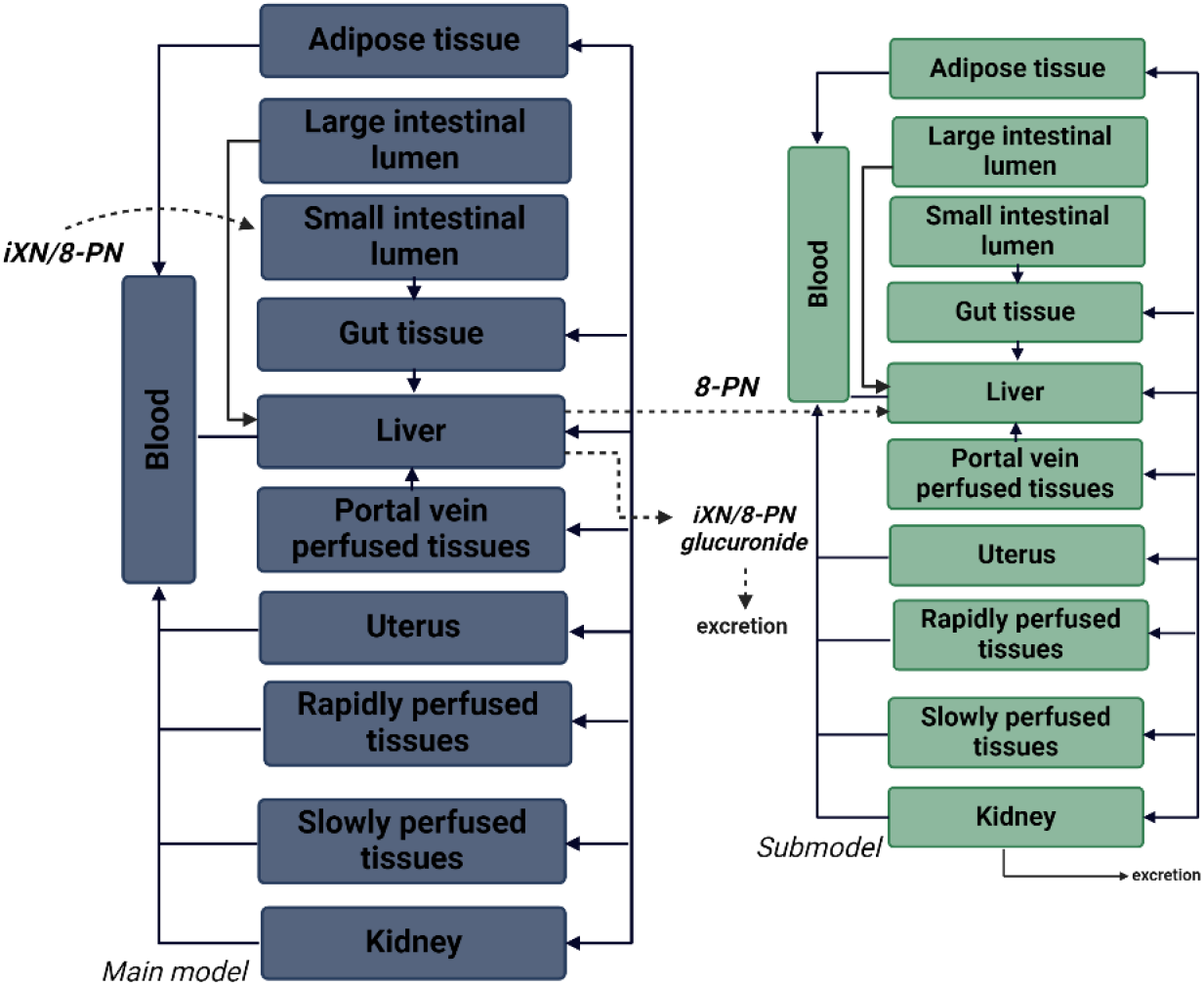
The structural concept of the PBK model, with the main model (blue) for iXN and the submodel (green) for 8-PN. The submodels for iXN and 8-PN glucuronides are not shown in the model. The dashed lines indicate transferring processes between the two submodels, uptake or excretion.

### 2.5. Cell culture and Caco-2 permeability assay

The Caco-2 human colon carcinoma cell line was cultured in Dulbecco’s Modified Eagle Medium (DMEM) supplemented with 1% (v/v) fetal bovine serum (FBS) and 1% (v/v) penicillin/streptomycin (P/S) for three weeks before seeding into Transwell® 0.4 µm pore polycarbonate membrane 12-well plates. The media was changed every second day for 24 ± 2 days prior to performing permeability assays. The cells were seeded on Transwell® polycarbonate membrane and the permeability assays were performed on day 24 or ± 2 days. The cell culture media was changed 12-24 h prior to performing the permeability assays. Additionally, the transepithelial electrical resistance (TEER) of each monolayer was measured and was well above 300 Ωcm^-2^, indicating good monolayer integrity. On the day of the permeability assays, the Caco-2 monolayer was washed with HBSS containing 25 mM HEPES (HBSS/HEPES) at pH 7.4 on both apical and basolateral side for 20 min on a shaking incubator (50 rpm, 37 °C). Test solutions of 50 µM iXN and 8-PN in HBSS/HEPES were prepared from stock solutions in DMSO (DMSO concentration was below 0.1%). The test solutions of iXN and 8-PN were added to the apical side of the monolayers and blank HBSS/HEPES to the basal side for measuring the apical to basolateral side permeability. From the basolateral side, samples were collected at selected time points (0, 5, 10, 20, 30, 60, 90, 120 min) and the apical side at time points 0 and 120 min. The amount of sample taken from the basolateral site was always replaced with the same amount of HBSS/HEPES. All samples were analyzed by LC-MS. Afterwards, the Caco-2 cell monolayer was washed three times with ice-cold PBS, the cells were detached by scraping, and ice-cold methanol was added to collect them for LC-MS analysis. Four independent experiments were performed, each comprised of four technical replicates.

### 2.6. Model parametrization

The PBK model was parameterized using a standard average female body weight of 60 kg. Relative tissue volumes and blood flow rates to the respective organs reported for women (24), small and large intestinal lumen volumes (25), and gastrointestinal transit times (26) were incorporated. The relative tissue volume and blood flow rates for the portal vein-perfused tissues were calculated as a single value. Gastrointestinal anatomical parameters were calculated as previously described (25). The glomerular filtration rate was obtained from the literature and set to 125 mL/min/(173 m^2^)(27). Compound-specific physiochemical parameters were calculated from LogP and pK_a_ values for iXN (LogP 4.1, pK_a_ 7.6 (28)), 8-PN (LogP 4.3, pK_a_ 7.7 (28)) and their glucuronides (iXNGluc, LogP 1.61, pK_a_ 8.55; 8-PNGluc LogP 1.34, pK_a_ 7.87 obtained from ChemDraw 19.0 Software) by using the QIVIVE toolbox (29). The toolbox calculates the tissue partition coefficient on the Rodges and Rowland method and the fraction unbound in plasma (fup) on Lobell and Sivarajah (30, 31). The K_m_ (17.17 µM) and V_max_ (5.47 pmol/min/pmol) for the conversion of iXN to 8-PN in the liver were reported in the literature, measured in human liver microsomes by CYP1A2 (11). The CYP1A2 conversion of iXN to 8-PN was scaled based on the intersystem extrapolation factor for CYP1A2 (ISEF), the P450 abundance pmol/mg protein, and the amount of microsomes/ liver weight (32). The absorption of iXN and 8-PN was predicted based on the apparent permeability coefficient (P_app_) measured *in vitro* and scalled to an *in vivo* P_app_ as described below. All parameters used can be found in the PBK model code in the supporting information. The input dose was based on a human intervention study in which postmenopausal women took three doses of a hops supplement (33). The data measured in the study was used to evaluate the accuracy of the PBK model predictions.

### 2.7. Sensitivity analysis

A sensitivity analysis was performed to evaluate the effect of parameter variation on the model predictions. To perform the sensitivity analysis, each parameter was increased by 5%, and the model was run at the dose given in the human intervention study. Using the formula developed by Evans and Andersen (34):

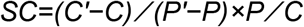

The normalized sensitivity coefficients (SC) were calculated, with C and C’ referring to the peak concentration of iXN in blood with unchanged or increased parameters, respectively, P and P’ referring to the value of the unchanged or increased parameter of interest.

### 2.8. Predictions of iXN and 8-N concentrations in tissues of interest

The PBK model was used to predict systemic and tissue concentrations of both iXN and 8-PN in all tissue compartments, including the uterus, which is considered of high relevance. The intake was based on different dosing scenarios administered *in vivo* in the study by van Breemen *et al*. (33).

### 2.9. LC-MS analysis of permeability assays and calculation of absorption kinetics

Samples from the Caco-2 permeability studies were centrifuged at 20’ 817 g for 15 min at 4 °C prior to LC-MS analysis. All samples were analyzed using a Vanquish HPLC (Thermo Fischer Scientific) coupled to an IDX tribrid mass spectrometer (Thermo Scientific). The chromatographic separation of analytes was achieved using a Phenomenex Synergi™ 4 µm Polar-RP (80 Å, 30 × 2 mm) column. The column compartment was set to 40 °C and the samples were stored in the autosampler cooled to 4 °C. The injection volume was 5 µL. The mobile phases were water (solvent A), and acetonitrile (solvent B) LC-MS grade, each supplemented with 0.1% formic acid. The flow rate was 0.7 mL/min, and the gradient was as follows: 0 min (5% B), 0.1 min (5% B), 1.8 min (100% B), 2.4 min (5% B), and 2.7 min (5% B). To protect the MS from HBSS, for the first 1.25 and last 0.6 min, the divert valve was set to waste. The MS parameters were negative ion spray voltage at 2500 V, sheath gas, auxiliary gas, and sweep gas at 60, 15, and 2 Arb, respectively, and ion transfer tube and vaporizer temperature at 350 °C and 400 °C, respectively. The P_app_ was calculated using the following formula (35):

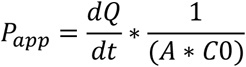

In the formula, dQ/dt is the steady-state flux (µM/s), change of concentration over time, A is the surface area of the Transwell®, and C0 is the initial concentration (µM) in the apical side. The mean P_app_ values for iXN and 8-PN derived from four independent experiments were scaled to *in vivo* P_eff_ (effective permeability) values using the algorithm of Sun *et al*. (36). These values were implemented in the PBK model in order to predict the absorption of both iXN and 8-PN absorption.

## 3. Results

### 3.1. *In vitro* glucuronide conjugation of iXN and 8-PN

Kinetic parameters for liver and intestinal phase II rates of glucuronidation of iXN and 8-PN were determined by incubation with human pooled hepatic and intestinal S9 fractions. For both iXN and 8-PN, the formation of glucuronides was observed as the appearance of one peak that was observed only when incubations were performed in the presence of UDPGA. No peaks corresponding to glucuronides were observed in the respective control without UDPGA. The glucuronide formation followed Michaelis-Menten kinetics, resulting in affinity constants (K_m_) of 6.33 µM for hepatic and 6.06 µM for intestinal S9 fraction for iXN. The V_max_ values for iXN were equal to 1.6 and 0.79 nmol/mg/min for hepatic and intestinal S9 fractions, respectively (Figure 3). The K_m_ values of 8-PN were 0.79 µM and 0.32 µM and V_max_ 0.76 and 1.250 nmol/mg/min for hepatic and intestinal S9, respectively. These results suggest that the glucuronidation of both iXN and 8-PN occurs rapidly in both, hepatic and intestinal tissues.

**Figure 3.**
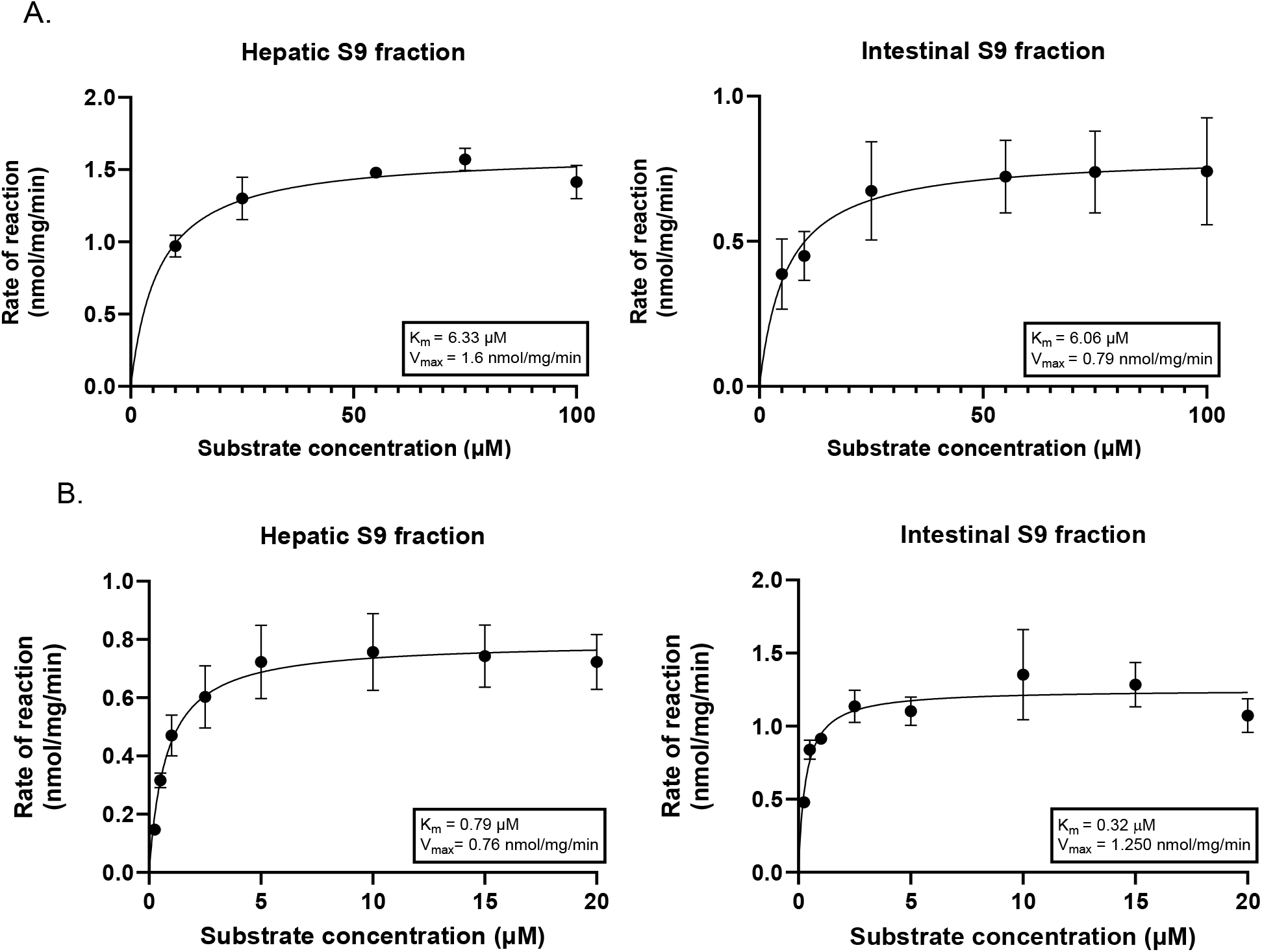
*In vitro* glucuronidation rates of iXN (A) and 8-PN (B) by hepatic and small intestinal S9 fractions. The iXN/8-PN-concentration-dependent formation rates are shown. Data are shown as mean ± SD of three experiments. The lines represent curve fits to the Michaelis-Menten equation.

### 3.2. Caco-2 cell permeability assay

To estimate the gastrointestinal permeability of iXN and 8-PN, Caco-2 (human colorectal adenocarcinoma) cells were cultured to form a monolayer in a Transwell® polycarbonate membrane system. The transfer of the compounds from the apical to the basolateral side was measured. Concentrations of both substances increased linearly in the basolateral compartment over at least 90 min. The P_app_ was calculated to be 6.06 ± 2.98 × 10^−6^ cm/sec for iXN, and 7.22 ± 1.66 × 10^−6^ cm/sec for 8-PN, respectively. These values indicate moderate permeability for both compounds, with 8-PN transferring slightly faster than iXN.

### 3.3. Comparison of the PBK model predictions with *in vivo* data from a human intervention study

The fully parameterized PBK model was run using starting doses of 0.8 mg, 1.6 mg, and 3.2 mg iXN, matching the conditions of a human intervention study (33). Serum concentrations (C_max_ values) of total iXN (iXN + iXNGluc) were predicted and compared to the previously published *in vivo* results (Figure 4A). These values matched well, supporting the accuracy of the model. The time required to reach the C_max_ (T_max_) predicted by the model was 7 h 47 min for the lowest and mid-dose and 7 h 45 min for the highest administered dose. In the dietary intervention study, the reported time required to reach the maximal concentration in blood ranged from 1.8 h up to 7.5 h across the different doses, indicating a large deviation amongst the subjects (33).

**Figure 4.**
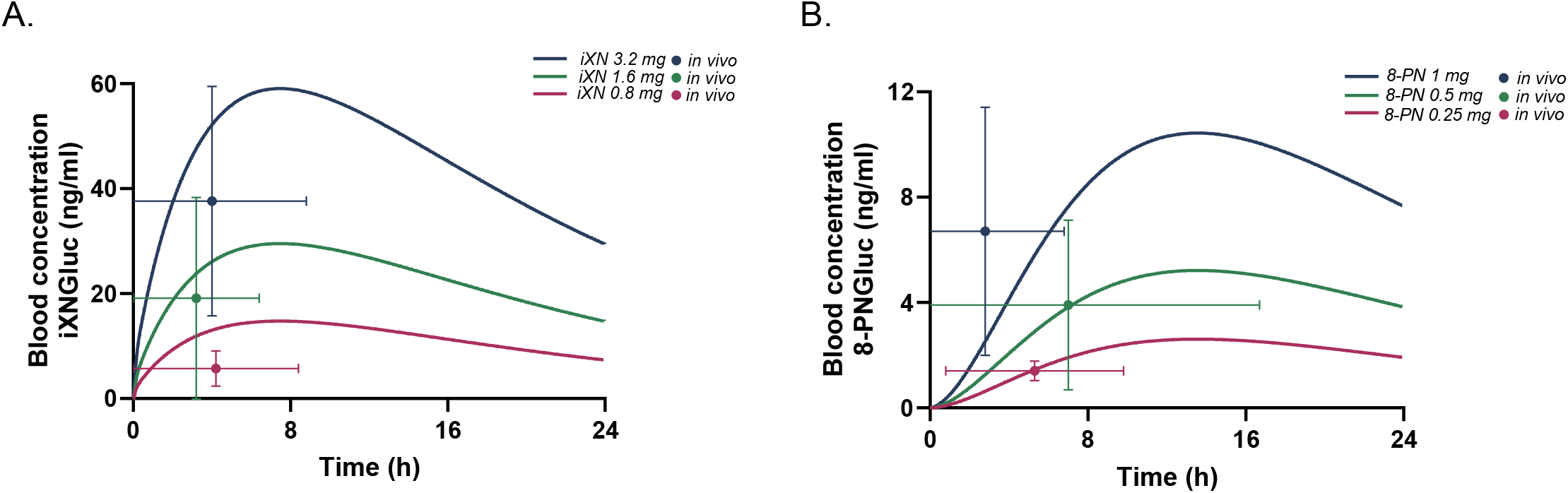
Curves representing predicted blood concentrations of iXN glucuronide (A) and 8-PN glucuronide (B) after supplementation with three doses of iXN and 8-PN from a hop supplement, compared with previously reported blood C_max_ and T_max_ values by van Bremen *et al*. (2014) (33). Curves represent PBK model–simulated blood concentrations over time, while circles represent the mean ± SD of C_max_ and T_max_ reported in the human intervention study.

Considering the large variation detected in the samples of the selected human intervention study, the model’s predictions were considered sufficiently accurate, and the small deviation in T_max_ values as compared to reported average values was considered acceptable. Additionally, the simulated C_max_ values for 8-PN glucuronide were compared to measured 8-PN blood concentrations (Figure 4B). These closely matched all three *in vivo* doses. Taking all comparisons of predicted to measured C_max_ values into consideration, the model provided adequate accuracy in predicting time-dependent blood concentrations of iXN and 8-PN glucuronides.

### 3.4. Sensitivity analysis

A sensitivity analysis was performed to assess which physiological parameters highly influence predicted blood concentrations of iXN and iXNGluc (Figure 5A). Exposure to a single dose of 0.8 mg iXN was modeled, corresponding to the lowest used dose by van Bremen *et al. (33)*. The predicted blood concentrations of iXN and iXNGluc were predominantly affected by body weight and the logP_app_ value of iXN. Parameters describing hepatic and intestinal metabolism of iXN also affected systematically available concentrations substantially. Specifically, the hepatic metabolism had a considerable impact with a sensitivity coefficient (SC) of -0.92 for V_max_ and 0.97 for K_m_. Besides metabolism, gastrointestinal parameters, including passing time through the small intestine and the area of small and large intestines, had SCs of -0.44, -0.16, and 0.57, respectively. Additionally, iXNGluc levels were slightly affected by the parameters describing glomerular filtration rate (GFR), slowly perfused tissue/blood partition coefficient (PSIXNGluc), fat/blood partition coefficient (PFIXNGluc), and faction unbound in plasma (FUIXNGluc). To a lesser extent, iXNGluc concentrations were influenced by gastrointestinal physiological parameters, including the volume and area of the small and large intestines. The predictions were also affected by the physiological parameter for the relative volume of adipose tissue. Regarding 8-PN, the sensitivity analysis was performed the same way as for iXN using exposure to a single dose of 0.25 mg 8-PN corresponding to the lowest dose by van Bremen *et al*. (33) (Figure 5B). The predicted blood concentrations of 8-PN and 8-PNGluc were affected predominantly by the same parameters as for iXN and iXNGluc including the logP_app_ value of 8-PN being the most influential parameter.

**Figure 5.**
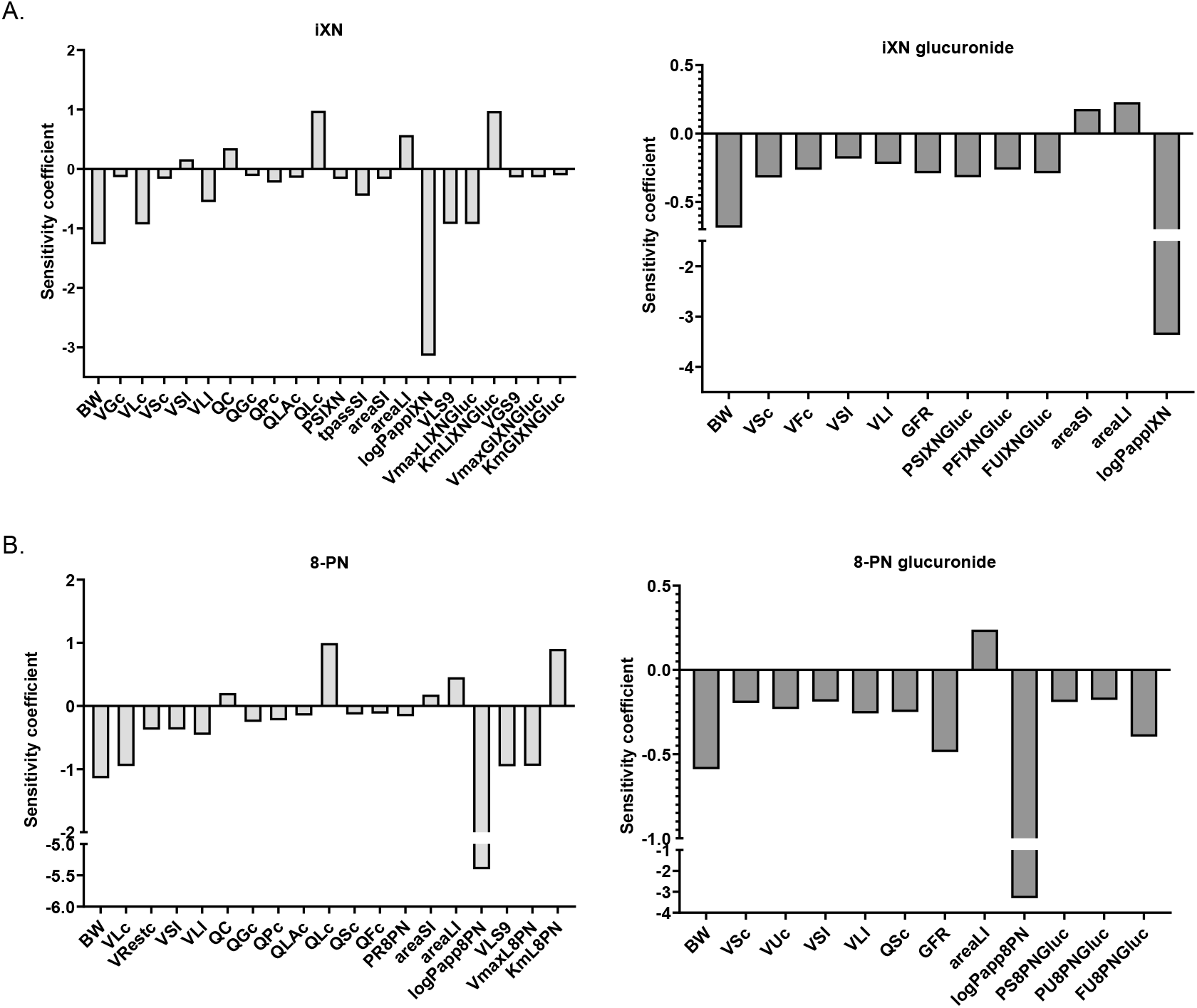
Sensitivity of model parameters (A. iXN and iXNGluc, B. 8-PN and 8-PNGluc), displayed as normalized sensitivity coefficients (SC) that were calculated from the alteration of blood C_max_ values caused by a 5% elevation of the respective parameter. Parameters with an absolute SC > 0.1 are shown and abbreviated as: BW, bodyweight; VGc, relative gut tissue (small intestine) volume; VLc, relative liver tissue volume; VSc, relative slowly perfused tissues volume; VFc, relative adipose tissue volume; VUc, relative tissue volume of the uterus; Vrestc, relative tissue volumes of all tissue not included in the model; VSI, volume of the small intestinal lumen; VLI, volume of the large intestinal lumen; QC, cardiac output; QGc, fraction of blood flow to gut; QPc fraction of blood flow to portal vein perfused tissues; QLA, fraction of blood flow to liver via artery; PSIXN/PSIXNGluc/ PS8PNGluc, slowly perfused tissue/blood partition coefficient; tpassSI, passing time through the small intestine; areaSI, surface area of the small intestinal lumen; areaLI surface area of large intestinal lumen; logPappIXN, logarithm of the permeability coefficient across the intestinal barrierof iXN; logPapp8PN, logarithm of the permeability coefficient across the intestinal barrier of 8-PN; VLS9, scaling factor for S9 protein to liver tissue; VmaxLIXNGluc, maximal velocity of hepatic glucuronidation of iXN, KmLIXNGluc, affinity constant for hepatic iXN glucuronidation; VGS9, scaling factor for S9 protein to intestinal tissue; VmaxGIXNGluc, maximal velocity of intestinal glucuronidation of iXN, KmGIXNGluc, affinity constant for intestinal iXN glucuronidation; GFR, glomerular filtration rate; PFIXNGluc, fat/blood partition coefficient; FUIXNGluc, FU8PNGluc faction unbound of iXNGluc and 8-PNGluc; PR8PN, rapidly perfused tissue/blood partition coefficient; QSc, fraction of blood flow to slowly perfused tissues; QFc, fraction of blood flow to fat; VmaxL8PN, maximal velocity of hepatic 8-PN glucuronidation; KmL8PN, affnity constant for hepatic 8-PN glucuronidation.

### 3.5 Predictions of iXN and 8-PN concentrations in blood and tissues of interest

Following the model evaluation, the model was used to simulate the systemic concentrations of iXN and 8-PN in their aglyconic bioactive form, based on three different dosing scenarios described in a human intervention study (33). Following the highest dose (3.6 mg of iXN and 1 mg of 8-PN daily), the maximal blood levels of iXN and 8-PN were predicted to be 0.13 nM and 0.008 nM, respectively (Figure 6). Additionally, following the same scenario, tissue levels of iXN and 8-PN were predicted to be in the low picomolar range for all selected organs of interest (i.e. liver, kidneys, uterus, adipose tissue; Figure 7). Of note, both for iXN and 8-PN our model predicts the substances to be slightly accumulated in adipose tissue after 24 h.

**Figure 6.**
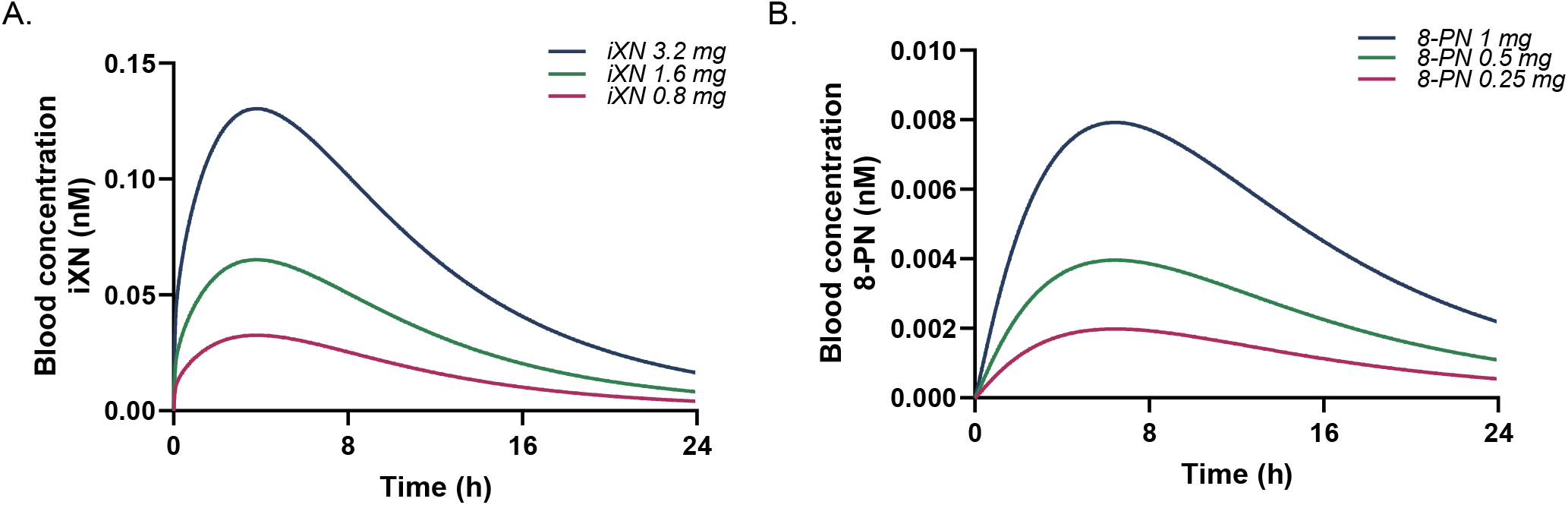
Blood concentrations of iXN (A) and 8-PN (B) following three different dosing scenarios after supplementation with a hop supplement, as predicted by the PBK model. Blue lines represent the highest dose, green middle dose and pink represents the lowest administered dose.

**Figure 7.**
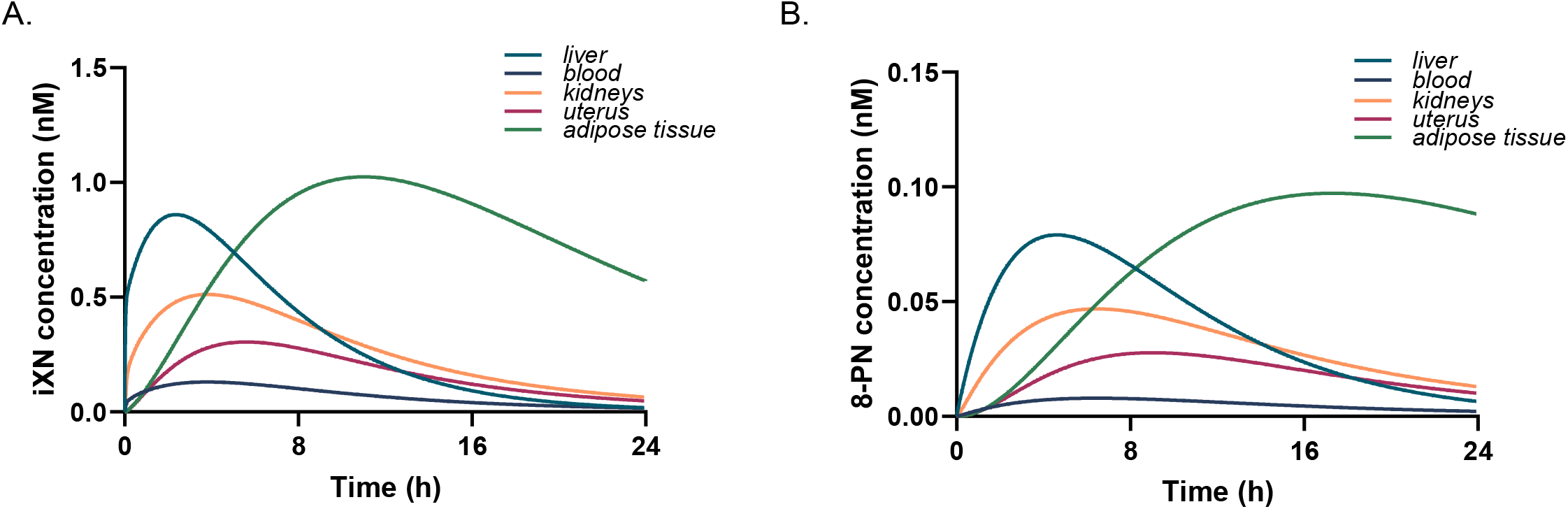
Predicted levels of iXN (A) and 8-PN (B) in organs of interest after supplementation with the highest dose of a hop supplement (3.2 mg of iXN and 1 mg of 8-PN).

Further, the model was used to predict blood and uterus concentrations of iXN and 8-PN following a dosing scenario of 1 beer daily for 7 days (Table 1). The specific amounts of iXN and 8-PN in beers used as an input dose for the PBK model were obtained from Stevens *et al*. (37) are shown in the supporting information (Table S1).

**Table 1.** Blood and uterus C_max_ of iXN and 8-PN after one beer a day for seven days scenario of highly and low-hopped beers.

| **Highly hopped beers** | **$C_{\max}$ blood iXN (pM)** | **$C_{\max}$ blood 8-PN (pM)** | **$C_{\max}$ uterus iXN (pM)** | **$C_{\max}$ uterus 8-PN (pM)** |
| --- | --- | --- | --- | --- |
| American porter | 59.32 | 2.32 | 139.13 | 8.19 |
| Strong ale | 153.20 | 1.11 | 360.10 | 3.94 |
| **Low hopped beers** |  |  |  |  |
| European lager | 1.77 | 0.01 | 4.16 | 0.04 |
| European pilsner | 25.43 | 0.21 | 59.63 | 0.74 |
| Alcohol-free beer | 4.90 | 0.03 | 11.49 | 0.11 |

## 4. Discussion

In this study, a PBK model was developed to estimate systemic concentrations of the estrogenic hop polyphenols isoxanthohumol and 8-prenylnaringenin, and their metabolites in women. The model integrates hepatic and intestinal glucuronidation kinetics measured using S9 fractions, human physiological parameters derived from the literature, tissue partitioning, and plasma binding estimated using quantitative structure-activity relationship (QSAR) tools. Initially, parameters for gastrointestinal permeability were taken from literature, but after sensitivity analysis indicated that the P_app_ of iXN and 8-PN was the most influential parameter for the prediction of C_max_, we decided to confirm these values experimentally. Therefore, *in vitro* measurements of intestinal permeability of iXN and 8-PN were conducted using a Caco-2 cell monolayer Transwell® system. P_app_ coefficients for iXN and 8-PN indicated a moderate permeability for both compounds, with values between those reported in the literature (38, 39). Variability of Caco-2 data, often observed across different laboratories, is attributed to factors including the passage number of the Caco-2 cells, pH conditions, and the integrity of the monolayer (40).

To evaluate the accuracy of the model’s predictions, the predicted blood concentrations of iXNGluc and 8-PNGluc were compared to the respective blood concentrations reported from a human intervention study (41). This comparison revealed only a 1.5-fold difference between predicted and *in vivo* measured levels of total (conjugated plus aglycone) iXN blood levels, whereas the predicted levels of 8-PN were even more closely aligned with those measured in the *in vivo* study (Figure 4). A limitation in evaluating the model was the limited availability of suitable human intervention studies. Research studies have primarily focused on xanthohumol, the precursor of iXN, administered as a single compound in rats (42) and humans (43) to monitor the *in vivo* formation of iXN and 8-PN. We identified three studies that administered iXN (33, 44, 45), but also administered the compound in mixtures with co-occurring polyphenols, including 8-PN, at different doses. Therefore, the blood levels of 8-PN could result from both, the initial dose of 8-PN and/or its conversion from iXN. From the published human intervention studies, the study by van Breemen *et al. (33)* was selected for model evaluation because of its detailed reporting of hop polyphenol content in the supplements and pharmacokinetic parameters, including C_max_ and T_max_ for both iXN and 8-PN. In their study, postmenopausal women were taking a hop supplement containing XN, iXN, 8-PN, and 6-prenylnaringenin either at low, medium, or high doses for 5 days. The applicability of their study to evaluate PBK models is limited by the small sample size (5 subjects), a treatment period of 5 days per dose, and the absence of additional information, e.g. reporting of individual values or subject physiology (BW, BMI). The study reported two peak concentrations of iXN and 8-PN in blood, indicating the occurrence of enterohepatic recirculation. Future models of hop polyphenols could consider incorporating enterohepatic circulation.

The PBK model was parameterized to incorporate the hepatic conversion of iXN to 8-PN by the human liver enzyme CYP1A2, based on *in vitro* studies with CYP1A2 and human microsomes (11). CYP1A2 activity is influenced by genetic polymorphisms and environmental exposures that act as inducers (e.g., cigarette smoking) or inhibitors (e.g., oral contraceptives)(46). These factors could impact systemic levels of 8-PN after iXN exposure, suggesting variable metabolic conversion among individuals. For future studies, it could be interesting to further extend the PBK model to account for genetic polymorphisms of CYP1A2, which would potentially predict the systemic concentrations of target compounds for each CYP1A2 genotype group.

Based on realistic dosing scenarios, the PBK model provides initial insights into tissue concentrations of both iXN and 8-PN, suggesting maximal concentrations in the low picomolar range. In a recent study, a detailed assessment of 8-PN was performed by comparing various *in vitro* and *in vivo* methods standardized by the OECD (47). The results of the study showed POD for 8-PN at 0.29 μM and 0.029 μM for ER-α dimerization and ER transactivation. Additionally, it was shown that 8-PN exhibits significant estrogenic activity, with PC50 values (corresponding to the dose required to cause 50% of the system’s maximal effect) of 2.14 μM for ER-α dimerization and 139 nM for ER transactivation. The ER-α dimerization for all compounds tested in the study was less sensitive compared to the ER transactivation. Moreover, a significant increase in uterine weight was observed in an *in vivo* uterotrophic assay, highlighting the estrogenic potential of 8-PN.

Even though this PBK model predicts significantly lower systemic concentrations at the used doses, higher 8-PN levels might be achieved via repeated exposure to hop supplements or hopped beverages. To test this, the model was also used to predict blood and uterus concentrations of 8-PN and iXN based on highly and low-hopped beer containing both iXN and 8-PN, simulating a scenario of one beer per day for seven days. The model predicted concentrations below 0.1 nM for both high and low-hopped beer scenarios. The internal levels are lower than after hops supplementation.

In an *in vitro* study using Ishikawa cells, slight synergistic estrogenic effects of iXN and 8-PN were demonstrated, indicating an enhancement of estrogenic activity upon the combination of these compounds (48). This interaction suggests that even at low concentrations, iXN and 8-PN could potentially influence estrogen receptor-mediated responses, highlighting the need for a thorough further evaluation. Additionally, the presence of endogenous estrogens *in vivo* adds another layer of complexity to their safety testing, as these hormones could potentially interact with the compounds being studied. It is crucial to recognize that the current model includes hepatic and intestinal tissue metabolism and, therefore, does not consider the potential conversion of iXN to 8-PN that was previously demonstrated to be catalyzed by the human gut microbiome. The respective O-demethylation was reported for specific bacteria occurring in the human gastrointestinal tract, e.g. for *Eubacterium limosum* and *Eubacterium rammulus* (49, 50). Additionally, a dietary intervention study of fifty healthy postmenopausal women who consumed a hop supplement showed the ability of the fecal microbiome to convert iXN to 8-PN with varying efficiency, resulting in a classification of poor, moderate, and strong 8-PN producers (51). This variability underscores the potential importance of microbial conversion and the pronounced interindividual differences in metabolism. The incorporation of gut microbial iXN to 8-PN conversion as a separate metabolic compartment in the PBK model could be a valuable approach to differentiate between different metabolizing groups and is envisioned for a follow-up study.

## 5. Conclusion

The PBK model estimates the systemic and tissue concentrations of hop-derived iXN and 8-PN upon different dosing scenarios. Both iXN and 8-PN blood levels were estimated at low picomolar ranges upon different dosing scenarios. The model serves as a tool for future quantitative *in vitro* to *in vivo* extrapolation (QIVIVE) to assess the likelihood of effects occurring *in vivo*.

## Supporting information

Supplementary model code

## Acknowledgments

This research was funded by Swiss Center for Applied Human Toxicology (SCAHT). Open access funding was provided by ETH Zurich. GA was also supported with a fellowship from the Future Food Initiative, a program run by the World Food System Center of ETH Zurich, the Integrative Food and Nutrition Center of EPFL and their industrial partners. The Dutch Ministry of Agriculture, Fisheries, Food Security and Nature (Ministerie van Landbouw, Visserij, Voedselzekerheid en Natuur) is gratefully acknowledged for providing financial support (KB-37-002-023). The authors are grateful to Amrei Rolof for helping with the 8-PN glucuronidation experiments.

## Declaration of Generative AI and AI-assisted technologies in the writing process

During the preparation of this work the authors used DeepL, Grammarly and ChatGPT 4 in order to refine and detect grammar and spelling errors. After using these tools, the authors reviewed and edited the content as needed and take full responsibility for the content of the publication.

## Author Contributions

**Maja Stevanoska**: Conceptualization, Methodology, Visualization, Investigation, Writing - Original Draft, Writing - Review & Editing; **Karsten Beekmann** Supervision, Methodology, Writing - Review & Editing; **Ans Punt:** Supervision, Methodology; **Shana J. Sturla:** Supervision, Writing - Review & Editing. **Georg Aichinger**: Supervision, Conceptualization, Methodology, Writing - Review & Editing;

