## Supplementary model code for "Predicting *in vivo* concentrations of dietary hop phytoestrogens by physiologically based kinetic modeling"

;Date: 2024-06-06
;Developed by: Maja Stevanoska, Georg Aichinger
;Compiled by: Maja Stevanoska
;Organization: Laboratory of Toxicology, D-HEST, ETH Zurich
;To execute the model, copy-paste the text to Berkeley Madonna

;Model code:

;=====================================================================

{Globals}

;=====================================================================

;Physiological parameters

;=====================================================================

;bodyweight

BW = 60	;kg	;BW of mean Caucasian female	;reference: (1)

;--------------------------------------------------------------------------------------------------------------------

;relative tissue volumes ;fraction of BW ;reference: (1)

VGc = 0.014666	;fraction of gut (small intestine) tissue

VLc = 0.02333	;fraction of liver tissue

VPc = 0.0216666	;fraction portal vein perfused tissue (stomach, spleen, pancreas, large intestine)

VRc = 0.033283	;fraction of rapidly perfused tissue (heart, lungs, brain)

VSc = 0.456666	;fraction of slowly perfused tissue (bone, skin, muscle)

VFc = 0.31666	;fraction of fat (adipose) tissue

VKc = 0.004583	;fraction of kidneys

VUc= 0.001333	;fraction of uterus

VBc = 0.068333	;fraction of blood

VRestc= 0.0594	;fraction of the rest of the body

;--------------------------------------------------------------------------------------------------------------------

;calculated tissue volumes

VG = VGc*BW	;L or Kg ;volume of gut tissue (calculated)

VL = VLc*BW	;L or Kg ;volume of liver tissue (calculated)

VP = VPc*BW	;L or kg ;volume of portal vein perfused tissue (calculated)

VR = VRc*BW	;L or Kg ;volume of rapidly perfused tissue (calculated)

VS = VSc*BW	;L or Kg ;volume of slowly perfused tissue (calculated)

VF = VFc*BW	;L or Kg ;volume of fat tissue (calculated)

VK = VKc*BW	;L or Kg ;volume of kidney (calculated)

VB = VBc*BW	;L or Kg ;volume of blood (calculated)

VU= VUc*BW	;L or kg ;volume of uterus (calculated)

VSI = 9.03	;L	;volume of small intestine lumen	;reference: (2)

VLI = 7.5	;L	;volume of large intestine lumen	;reference: (2)

;--------------------------------------------------------------------------------------------------------------------

;Blood flow rates, human female	;reference: (1)

QC = 354	;L/h	;cardiac output, resting human female	;reference: (1)

QGc = 0.11	;fraction of blood flow to gut (small intestine)

QPc = 0.1	;fraction of blood flow to Portal vein perfused tissues

QLAc = 0.065	;fraction of blood flow to liver via artery

QLc = 0.27	;fraction of blood flow to liver in total

QRc = 0.195	;fraction of blood flow to rapidly perfused tissue (heart, lungs, brain)

QSc = 0.22	;fraction of blood flow to slowly perfused tissue (skin, bone, muscle)

QFc = 0.085	;fraction of blood flow to fat

QKc = 0.17	;fraction of blood flow to kidneys

QUc= 0.0004	;fraction of blood to uterus

QG = QGc*QC	;blood flow to gut (calculated)

QP = QPc*QC	;blood flow to portal vein perfused tissue (calculated)

QLA = QLAc*QC	;blood flow to liver via artery (calculated)

QL = QLc*QC	;blood flow to liver total (calculated)

QR = QRc*QC	;blood flow to rapidly perfused tissue (calculated)

QS = QSc*QC	;blood flow to slowly perfused tissue (calculated)

QF = QFc*QC	;blood flow to fat tissue (calculated)

QK = QKc*QC	;blood flow to kidney (calculated)

QU= QUc* QC	;blood flow to uterus (calculated)

;--------------------------------------------------------------------------------------------------------------------

;glomerular filtration rate	;reference: (3)

GFR = 125	;mL/min/173m2

;=====================================================================

;Physicochemical parameters

;=====================================================================

;partition coefficients ;calculated by Rodgers and Rowland method, qivivetools.wu.nl	;reference: (4)

;iXN in main model ;reference: (5)

PGIXN = 8.35	;gut/blood partition coefficient

PPIXN = 2.12	;portal vein perfused tissue/blood partition coefficient (spleen)

PLIXN = 4.19	;liver/blood partition coefficient

PRIXN = 5.52	;rapidly perfused tissue/blood partition coefficient	;weighted average

PSIXN = 6.2	;slowly perfused tissue/blood partition coefficient	;weighted average

PFIXN = 13.93	;fat/blood partition coefficient

PKIXN = 3.94	;kidney/blood partition coefficient

PUIXN= 2.48	;uterus/blood partition coefficient	;same as muscle

FUIXN = 0.026	;faction unbound plasma	;Lobell&Sivarajah algorithm	;reference: (4)

;iXN Glucuronide (IXNGluc) in sub-model

PGIXNGluc = 0.93 ;gut/blood partition coefficient

PPIXNGluc = 0.40 ;portal vein perfused tissue/blood partition coefficient ;spleen

PLIXNGluc = 0.57 ;liver/blood partition coefficient

PRIXNGluc = 0.68 ;rapidly perfused tissue/blood partition coefficient	;weighted average

PSIXNGluc = 0.72 ;slowly perfused tissue/blood partition coefficient	;weighted average

PFIXNGluc = 0.84 ;fat/blood partition coefficient

PKIXNGluc = 0.59 ;kidney/blood partition coefficient

PUIXNGluc= 0.46 ;uterus/blood partition coefficient	;same as muscle

FUIXNGluc = 0.363 ;faction unbound plasma	;Lobell&Sivarajah algorithm	;reference: (4)

;8PN in sub-model (LogP 4.3 pKa 7.7)	;reference: (5)

PG8PN = 12.62	;gut/blood partition coefficient

PP8PN = 3.18	;portal vein perfused tissue/blood partition coefficient (spleen)

PL8PN = 6.33	;liver/blood partition coefficient

PR8PN = 8.32	;rapidly perfused tissue/blood partition coefficient	;weighted average

PS8PN = 9.36	;slowly perfused tissue/blood partition coefficient	;weighted average

PF8PN = 24.25	;fat/blood partition coefficient

PK8PN = 5.93	;kidney/blood partition coefficient

PU8PN= 3.74	;uterus/blood partition coefficient same as muscle value

FU8PN = 0.023	;faction unbound plasma according to Lobell&Sivarajah ;reference: (4)

;8PNGluc in sub-model

PG8PNGluc = 0.54	;gut/blood partition coefficient

PP8PNGluc = 0.28	;portal vein perfused tissue/blood partition coefficient

PL8PNGluc = 0.35	;liver/blood partition coefficient

PR8PNGluc = 0.42	;rapidly perfused tissue/blood partition coefficient ;weighted average

PS8PNGluc = 0.43	;slowly perfused tissue/blood partition coefficient ;weighted average

PF8PNGluc = 0.29	;fat/blood partition coefficient

PK8PNGluc = 0.39	;kidney/blood partition coefficient

PU8PNGluc= 0.31	;uterus/blood partition coefficient same as muscle value

FU8PNGluc = 0.323	;faction unbound plasma ;Lobell&Sivarajah algorithm	;reference: (4)

;--------------------------------------------------------------------------------------------------------------------

;molecular weight

MWIXN = 354.39	;molecular weight IXN

MWIXNGluc = 530.52	;molecular weight IXNGluc (obtained from ChemDraw 19.0 Software)

MW8PN= 340.375	;molecular weight 8PN

MW8PNGluc=516.5	;molecular weight 8PNGluc (obtained from ChemDraw 19.0 Software)

;--------------------------------------------------------------------------------------------------------------------

;Oral dose

ODOSEIXN=0.8	;mg	;oral dose iXN variable

AODOSEIXN=ODOSEIXN*1000/MWIXN	;µmol

ODOSE8PN= 0.25	;mg	;oral dose 8-PN variable

AODOSE8PN=ODOSE8PN*1000/MW8PN	;µmol

;--------------------------------------------------------------------------------------------------------------------

;gastrointestinal tract

tpassSI = 4.3	;h	;passing time through small intestine	;reference: (6)

Ksi = 1/tpassSI	;/h	;SI to LI transfer constant

tpassLI = 24.2	;h	;passing time through large intestine	;reference: (6)

Kli = 1/tpassLI	;/h	;LI to excretion transfer constant

;--------------------------------------------------------------------------------------------------------------------

;surface areas

areaSI = 72.1	;dm2 ;surface area of SI lumen ;reference: (2)

areaLI = 47.12	;dm2 ;surface area of LI lumen ;reference: (2)

;--------------------------------------------------------------------------------------------------------------------

;absorption/transfer rates iXN

logPappIXN = -5.217090041	;log(cm/s)	;measured Papp *in vitro*

PeffIXN = (3600/10)*(10^(0.6836*logPappIXN-0.5579))

;dm/h	;scaled according to Sun *et al*.	;reference: (7)

KaIXN = PeffIXN*areaSI	;L/h	;transfer rate of IXN from SI lumen to liver

KbIXN = PeffIXN*areaLI	;L/h	;transfer rate of IXN from LI lumen to liver

;absorption/transfer rates 8PN

logPapp8PN =-5.140978035	;log(cm/s)	;measured Papp in vitro ;log(cm/s)

Peff8PN = (3600/10)*(10^(0.6836*logPapp8PN-0.5579))

;dm/h	;scaled according to Sun *et al.*	;reference: (7)

Ka8PN = Peff8PN*areaSI	;L/h	;transfer rate of 8PN from SI lumen to liver

Kb8PN = Peff8PN*areaLI	;L/h	;transfer rate of 8PN from LI lumen to liver

;--------------------------------------------------------------------------------------------------------------------

;=====================================================================

;Kinetic parameters

;=====================================================================

;hepatic metabolism

;scaling factors

VLS9 = 107.3	;mg S9 protein/gram liver	;liver S9 protein yield	;reference: (8)

SFmprot=40	;(mg microsomal protein/g liver)	;liver microsome protein yield 	;reference: (9)

L=VLc*1000	;gram/kg BW	;fraction liver tissue

cCYP1A2 = 52	;(pmol/mg microsomal protein)	;CYP1A2 yield	;reference: (9)

ISEF= 0.17	;intersystem extrapolation factor for CYP1A2	;reference: (9)

;--------------------------------------------------------------------------------------------------------------------

;IXN phase II metabolism	;hepatic glucuronidation

VmaxLIXNGluc= 1.615 ;nmol/min/mg S9 protein ;unscaled maximum rate	;experimental

VmaxLIXNGlucs = VmaxLIXNGluc/1000*60* VLS9 *L*BW ;µmol/h	; scaled maximum rate

KmLIXNGluc = 6.33 ;µmol/L	;affinity constant	;experimental

;--------------------------------------------------------------------------------------------------------------------

;IXN phase I metabolism	;8PN formation in the liver

VmaxLiXN8PNc = 5.47	;(pmol/min)/pmol CYP1A2	;unscaled maximum rate ;reference: (10)

VmaxLiXN8PNs = (VmaxLiXN8PNc*cCYP1A2*SFmprot*ISEF)/1000000*60*L*BW

;µmol/h	;scaled maximum rate

KmLiXN8PN = 17.8	;µM	;affinity constant	;reference: (10)

FLiXN8PNc=VmaxLiXN8PNs*CVLIXN/(KmLiXN8PN+CVLIXN)	;function for hepatic 8PN formation

;--------------------------------------------------------------------------------------------------------------------

;intestinal metabolism

;scaling factors

VGS9 = 35.2	;mg S9 protein/gram SI ;SI S9 protein yield	;reference: (11)

G=VGc*1000	;g/kg BW	;fraction gut tissue

;--------------------------------------------------------------------------------------------------------------------

;IXN phase II metabolism	;intestinal glucuronidation

VmaxGIXNGluc = 0.7983	;nmol/min/mg S9 protein ;unscaled maximum rate	;experimental

VmaxGIXNGlucs = VmaxGIXNGluc/1000*60* VGS9 *G*BW ;µmol/h	;scaled maximum rate

KmGIXNGluc = 6.068	;µmol/L	;affinity constant	;experimental

;--------------------------------------------------------------------------------------------------------------------

;8PN phase II metabolism	;hepatic glucuronidation

VmaxL8PN = 0.7927 ;nmol/min/mg S9 protein	;unscaled maximum rate	;experimental

VmaxL8PNs = VmaxL8PN/1000*60*VLS9*L*BW	;µmol/h	;scaled maximum rate

KmL8PN = 0.7658	;µmol/L	;affinity constant	;experimental

;--------------------------------------------------------------------------------------------------------------------

;8PN phase II metabolism-glucuronidation intestinal tissue

VmaxG8PN = 1.25	;nmol*min-1*mg-1	;unscaled maximum rate	;experimental

VmaxG8PNs = VmaxG8PN/1000*60* VGS9 *G*BW	;µmol/h	;scaled maximum rate

KmG8PN = 0.3241	;µmol/L	;affinity constant	;experimental

;--------------------------------------------------------------------------------------------------------------------

;=====================================================================

;Main model calculations/dynamics: isoxanthohumol

;=====================================================================

;small intestine compartment

;ASIIXN: amount of IXN in small intestinal lumen ;µmol

CSIIXN=ASIIXN/VSI	;µmol/L

CSIIXNnm=CSIIXN*1000	;nM

;--------------------------------------------------------------------------------------------------------------------

;large intestinal lumen compartment

;ALIIXN: amount of IXN in large intestine lumen ;µmol

CLIIXN = ALIIXN/VLI	;µmol/Lmol

CLIIXNnm=CLIIXN*1000	;nM

;--------------------------------------------------------------------------------------------------------------------

;gut tissue compartment

;AGIXN: amount of IXN in gut tissue ;µmol

CGIXN = AGIXN/VG	;µmol/L

CVGIXN = CGIXN/PGIXN	;partition with gut tissue

CGIXNnm=CGIXN*1000	;nM

;--------------------------------------------------------------------------------------------------------------------

;liver compartment

;ALIXN: amount of IXN in liver	;µmol

CLIXN = ALIXN/VL	;µmol/L

CVLIXN = CLIXN/PLIXN	;partition with liver tissue

CLIXNnm=CLIXN*1000	;nM

;--------------------------------------------------------------------------------------------------------------------

;portal vein perfused tissue compartment

;APIXN: amount of IXN in spleen tissue ;µmol

CPIXN = APIXN/VP ;µmol/L

CVPIXN = CPIXN/PPIXN ;partition with portal vein perfused tissue

CPIXNnm=CPIXN*1000 ;nM

;----------------------------------------------------------------------------------------------------------------------

;blood compartment

CBIXN=ABIXN/VB ;µmol/L

CVBIXN = CBIXN*FUIXN ;free concentration in plasma

CBIXNug = CBIXN*MWIXN ;µg/L = ng/mL ;concentration in blood in ng/ml

CBIXNnm=CBIXN*1000 ;nM

;--------------------------------------------------------------------------------------------------------------------

;fat (adipose tissue) compartment

;AFXN: amount of IXN in fat ;µmol

CFIXN = AFIXN/VF	;µmol/L

CVFIXN = CFIXN/PFIXN	;partition with adipose tissue

CFIXNnm=CFIXN*1000	;nM

;--------------------------------------------------------------------------------------------------------------------

;rapidly perfused tissue

;ARIXN: amount of IXN in rapidly perfused tissue ;µmol

CRIXN = ARIXN/VR	;µmol/L

CVRIXN = CRIXN/PRIXN ;partition with rapidly perfused tissue

CRIXNnm=CRIXN*1000 ;nM

;--------------------------------------------------------------------------------------------------------------------

;slowly perfused tissue

;ASIXN: amount of IXN in slowly perfused tissue ;µmol

CSIXN = ASIXN/VS	;µmol/L

CVSIXN = CSIXN/PSIXN	;partition with slowly perfused tissue

CSIXNnm=CSIXN*1000	;nM

;--------------------------------------------------------------------------------------------------------------------

;kidney

;AKIXN: amount of IXN in kidney ;µmol

CKIXN = AKIXN/VK	;µmol/L

CVKIXN = CKIXN/PKIXN ;partition with kidney tissue

CKIXNnm=CKIXN*1000 ;nM

;--------------------------------------------------------------------------------------------------------------------

;uterus

;AUIXN: amount of iXN in uterus ;µmol

CUIXN= AUIXN/VU	;µmol/L

CUIXNnm= CUIXN*1000	;µmol/L

CVUIXN= CUIXN/PUIXN	;partition with uterus

CUIXNnm=CUIXN*1000	;nM

;--------------------------------------------------------------------------------------------------------------------

;=========================================================================

;Sub-model calculations/dynamics: iXN glucuronide

;=========================================================================

;--------------------------------------------------------------------------------------------------------------------

;gut tissue compartment

;AGIXNGluc: amount of IXN glucuronide in gut tissue ;µmol

CGIXNGluc = AGIXNGluc/VG	;µmol/L

CVGIXNGluc = CGIXNGluc/PGIXNGluc	;partition with gut tissue

;--------------------------------------------------------------------------------------------------------------------

;liver compartment

;ALIXNGluc: amount of IXN glucuronide in liver ;µmol

CLIXNGluc = ALIXNGluc/VL	;µmol/L

CVLIXNGluc = CLIXNGluc/PLIXNGluc	;partition with liver tissue

;--------------------------------------------------------------------------------------------------------------------

;fat (adipose tissue) compartment

;AFIXNGluc = amount of IXN glucuronide in fat tissue ;µmol

CFIXNGluc = AFIXNGluc/VF	;µmol/L

CVFIXNGluc = CFIXNGluc/PFIXNGluc	;partition with fat tissue

;--------------------------------------------------------------------------------------------------------------------

;portal vein perfused compartment

;APIXNGluc = amount of IXN glucuronide in portal vein perfused tissue ;µmol

CPIXNGluc = APIXNGluc/VP	;µmol/L

CVPIXNGluc = CPIXNGluc/PPIXNGluc	;partition with portal vein perfused tissue

;--------------------------------------------------------------------------------------------------------------------

;blood compartment

;ABIXNGluc: amount of IXN glucuronide in blood ;µmol

CBIXNGluc=ABIXNGluc/VB	;µmol/L

CVBIXNGluc = CBIXNGluc*FUIXNGluc	;concentration of free IXNGluc in plasma

CBIXNGlucnm=CBIXNGluc*1000

CBIXNGlucug = CBIXNGluc*MWIXNGluc ;concentration in blood in ng/mL

;--------------------------------------------------------------------------------------------------------------------

;rapidly perfused tissue

;ARIXNGluc = amount of iXN glucuronide in rapidly perfused tissue ;µmol

CRIXNGluc = ARIXNGluc/VR	;µmol/L

CVRIXNGluc = CRIXNGluc/PRIXNGluc	;partition with rapidly perfused tissue

;--------------------------------------------------------------------------------------------------------------------

;slowly perfused tissue

;ASIXNGluc = amount of iXN glucuronide in slowly perfused tissue ;µmol

CSIXNGluc = ASIXNGluc/VS	;µmol/L

CVSIXNGluc = CSIXNGluc/PSIXNGluc	;partition with slowly perfused tissue

;--------------------------------------------------------------------------------------------------------------------

;kidney

;AKIXNGluc: amount of IXN glucuronide in kidney ;µmol

CKIXNGluc = AKIXNGluc/VK	;µmol/L

CVKIXNGluc = CKIXNGluc/PKIXNGluc	;partition with kidney tissue

;--------------------------------------------------------------------------------------------------------------------

;uterus

;AUIXNGluc= amount of iXNGluc in uterus ;µmol

CUIXNGluc= AUIXNGluc/VU	;µmol/L

CUIXNGlucnm= CUIXNGluc*1000	;µmol/L

CVUIXNGluc= CUIXNGluc/PUIXNGluc	;partition with uterus

;--------------------------------------------------------------------------------------------------------------------

;=========================================================================

;Sub-model calculations/dynamics: 8PN

;=========================================================================

;--------------------------------------------------------------------------------------------------------------------

;liver compartment

;AL8PN: amount of 8PN in liver ;µmol

CL8PN= AL8PN/VL	;µmol/L

CVL8PN= CL8PN/PL8PN	;partition with liver tissue

CL8PNnm=CL8PN*1000	;nM

;--------------------------------------------------------------------------------------------------------------------

;gut tissue compartment

;AG8PN: amount of 8PN in gut tissue ;µmol

CG8PN= AG8PN/VG	;µmol/L

CVG8PN= CG8PN/PG8PN	;partition with gut tissue

CG8PNnm=CG8PN*1000	;nM

;--------------------------------------------------------------------------------------------------------------------

;blood compartment

;AB8PN: amount of 8PN in blood	;µmol

CB8PN=AB8PN/VB	;µmol/L

CVB8PN= CB8PN*FU8PN	;concentration of free 8PN in plasma

CB8PNnm=CB8PN*1000	;nM

CB8PNug = CB8PN*MW8PN	;concentration in blood in µg/L = ng/mL

;--------------------------------------------------------------------------------------------------------------------

;portal vein perfused compartment

;AP8PN: amount of 8PN in portal vein perfused tissue ;µmol

CP8PN= AP8PN/VP	;µmol/L

CVP8PN= CP8PN/PP8PN	;partition with portal vein perfused tissue

CP8PNnm=CP8PN*1000	;nM

;--------------------------------------------------------------------------------------------------------------------

;fat (adipose tissue) compartment

;AF8PN= amount of 8PN in fat tissue ;µmol

CF8PN= AF8PN/VF	;µmol/L

CVF8PN= CF8PN/PF8PN	;partition with fat tissue

CF8PNnm=CF8PN*1000	;nM

;--------------------------------------------------------------------------------------------------------------------

;rapidly perfused tissue

;AR8PN: amount of 8PN in rapidly perfused tissue ;µmol

CR8PN= AR8PN/VR	;µmol/L

CVR8PN= CR8PN/PR8PN	;partition with rapidly perfused tissue

;--------------------------------------------------------------------------------------------------------------------

;slowly perfused tissue

;AS8PN: amount of 8PN in slowly perfused tissue ;µmol

CS8PN= AS8PN/VS	;µmol/L

CVS8PN= CS8PN/PS8PN	;partition with slowly perfused tissue

CS8PNnm=CS8PN*1000	;nM

;--------------------------------------------------------------------------------------------------------------------

;kidney

;AK8PN: amount of 8PN in kidney ;µmol

CK8PN= AK8PN/VK	;µmol/L

CVK8PN= CK8PN/PK8PN	;partition with kidney tissue

CK8PNnm=CK8PN*1000	;nM

;--------------------------------------------------------------------------------------------------------------------

;large intestinal lumen compartment

;ALI8PN: amount of 8PN in large intestine lumen ;µmol

CLI8PN = ALI8PN/VLI	;µmol/L

CLI8PNnm=CLI8PN*1000	;nM

;--------------------------------------------------------------------------------------------------------------------

;small intestinal lumen compartment

;ASI8PN: amount of 8PN in large intestine lumen ;µmol

CSI8PN = ASI8PN/VSI	;µmol/L

CSI8PNnm=CSI8PN*1000	;nM

;--------------------------------------------------------------------------------------------------------------------

;uterus

;AU8PN: amount of 8PN in uterus ;µmol

CU8PN= AU8PN/VU	;µmol/L

CU8PNnm=CU8PN*1000	;nM

CVU8PN= CU8PN/PU8PN	;partition with uterus

;--------------------------------------------------------------------------------------------------------------------

;=========================================================================

;Sub-model calculations/dynamics: 8PNGluc

;=========================================================================

;--------------------------------------------------------------------------------------------------------------------

;liver compartment

;AL8PNGluc: amount of 8PNGluc in liver ;µmol

CL8PNGluc= AL8PNGluc/VL	;µmol/L

CVL8PNGluc= CL8PNGluc/PL8PNGluc	;partition with liver tissue

;--------------------------------------------------------------------------------------------------------------------

;gut tissue compartment

;AG8PNGluc: amount of 8PNGluc in gut tissue ;µmol

CG8PNGluc= AG8PNGluc/VG	;µmol/L

CVG8PNGluc= CG8PNGluc/PG8PNGluc ;partition with gut tissue

;--------------------------------------------------------------------------------------------------------------------

;blood compartment

;AB8PNGluc: amount of 8PNGluc in blood	;µmol

CB8PNGluc=AB8PNGluc/VB	;µmol/L

CVB8PNGluc= CB8PNGluc*FU8PNGluc	;concentration of free 8PNGluc in plasma

CB8PNGlucnm=CB8PNGluc*1000	;nM

CB8PNGlucug = CB8PNGluc*MW8PNGluc	;concentration in blood in ng/mL

;--------------------------------------------------------------------------------------------------------------------

;portal vein perfused compartment

;AP8PNGluc= amount of 8PNGluc in portal vein perfused tissue ;µmol

CP8PNGluc= AP8PNGluc/VP	;µmol/L

CVP8PNGluc= CP8PNGluc/PP8PNGluc	;partition with portal vein perfused tissue

;--------------------------------------------------------------------------------------------------------------------

;fat (adipose tissue) compartment

;AF8PNGluc= amount of 8PNGluc in fat tissue	;µmol

CF8PNGluc= AF8PNGluc/VF	;µmol/L

CVF8PNGluc= CF8PNGluc/PF8PNGluc	;partition with fat tissue

;--------------------------------------------------------------------------------------------------------------------

;rapidly perfused tissue

;AR8PNGluc= amount of 8PNGluc in rapidly perfused tissue ;µmol

CR8PNGluc= AR8PNGluc/VR	;µmol/L

CVR8PNGluc= CR8PNGluc/PR8PNGluc	;partition with rapidly perfused tissue

;--------------------------------------------------------------------------------------------------------------------

;slowly perfused tissue

;AS8PNGluc= amount of 8PNGluc in slowly perfused tissue ;µmol

CS8PNGluc= AS8PNGluc/VS	;µmol/L

CVS8PNGluc= CS8PNGluc/PS8PNGluc	;partition with slowly perfused tissue

;--------------------------------------------------------------------------------------------------------------------

;kidney

;AK8PNGluc: amount of 8PNGluc in kidney ;µmol

CK8PNGluc= AK8PNGluc/VK	;µmol/L

CVK8PNGluc= CK8PNGluc/PK8PNGluc	;partition with kidney tissue

;--------------------------------------------------------------------------------------------------------------------

;uterus

;AU8PNGluc: amount of 8PNGluc in uterus ;µmol

CU8PNGluc= AU8PNGluc/VU	;µmol/L

CU8PNGlucnm=CU8PNGluc*1000	;nM

CVU8PNGluc= CU8PNGluc/PU8PNGluc	;partition with slowly perfused tissue

;--------------------------------------------------------------------------------------------------------------------

;=====================================================================

;Model mass balance calculation

;=====================================================================

Total = AODOSEIXN + AODOSE8PN

Calculated=ASIIXN + ALIIXN + AGIXN+ ALIXN + APIXN + AFIXN + ABIXN+ ARIXN + ASIXN+ AKIXN+ ALIXNGluc + AGIXNGluc+ AL8PN+ IXNExUr + APIXNGluc + ABIXNGluc + AFIXNGluc+ ARIXNGluc + ASIXNGluc + AUIXNGluc+ AKIXNGluc + AG8PN + AB8PN+ AP8PN+ AR8PN + AS8PN + AK8PN+ AF8PN + ALI8PN +ASI8PN +AU8PN + Eliminate8PN+IXN8PNExUr+ IXNGlucExUr+ IXex + AL8PNGluc + AG8PNGluc+ AB8PNGluc+ AP8PNGluc+ AR8PNGluc + AS8PNGluc + AK8PNGluc+ AF8PNGluc +PNGlucEx +AU8PNGluc

ERROR= ((Total-Calculated)/Total + 1E-30)*100

MASSBAL=Total-Calculated + 1

{End Globals}

;=====================================================================

{Top model}

{Reservoirs}

d/dt (ASIIXN) = - transSI - transSItoG

INIT ASIIXN = AODOSEIXN

LIMIT ASIIXN >= 0

d/dt (ALIIXN) = + transSI - transLItoL - transLI

INIT ALIIXN = 0

LIMIT ALIIXN >= 0

d/dt (AGIXN) = + artG - venG + transSItoG - FGIXNGluc

INIT AGIXN = 0

LIMIT AGIXN >= 0

d/dt (ALIXN) = + transLItoL + venP + venG + artL - venL - FLIXNGluc - FLiXN8PNc

INIT ALIXN = 0

LIMIT ALIXN >= 0

d/dt (APIXN) = - venP + artP

INIT APIXN = 0

LIMIT APIXN >= 0

d/dt (ABIXN) = - artP - artG - artF + venF - artR + venR - artS + venS - artK + venK - artL + venL + venU - artU

INIT ABIXN = 0

LIMIT ABIXN >= 0

d/dt (AFIXN) = + artF - venF

INIT AFIXN = 0

LIMIT AFIXN >= 0

d/dt (ARIXN) = + artR - venR

INIT ARIXN = 0

LIMIT ARIXN >= 0

d/dt (ASIXN) = + artS - venS

INIT ASIXN = 0

LIMIT ASIXN >= 0

d/dt (AKIXN) = + artK - venK - urineiXN

INIT AKIXN = 0

LIMIT AKIXN >= 0

d/dt (IXNExUr) = + urineiXN

INIT IXNExUr = 0

LIMIT IXNExUr >= 0

d/dt (IXex) = + transLI

INIT IXex = 0

LIMIT IXex >= 0

d/dt (AUIXN) = - venU + artU

INIT AUIXN = 0

LIMIT AUIXN >= 0

{Flows}

transSI = ASIIXN*Ksi

transLItoL = KbIXN*CLIIXN

venP = QP*CVPIXN

artP = QP*CBIXN

artG = QG*CBIXN

venG = QG*CVGIXN

artF = QF*CBIXN

venF = QF*CVFIXN

artR = QR*CBIXN

venR = QR*CVRIXN

artS = QS*CBIXN

venS = QS*CVSIXN

artK = QK*CBIXN

venK = QK*CVKIXN

urineiXN = GFR*CVBIXN*60/1000

artL = QLA*CBIXN

venL = QL*CVLIXN

transSItoG = KaIXN*CSIIXN

transLI = ALIIXN*Kli

venU = QU*CVUIXN

artU = QU*CBIXN

{Submodel "IXNglucuronide"}

{Reservoirs}

d/dt (ALIXNGluc) = + FLIXNGluc + venP1 + venG1 - venL1 + artL1

INIT ALIXNGluc = 0

LIMIT ALIXNGluc >= 0

d/dt (AGIXNGluc) = + FGIXNGluc - venG1 + artG1

INIT AGIXNGluc = 0

LIMIT AGIXNGluc >= 0

d/dt (APIXNGluc) = + artP1 - venP1

INIT APIXNGluc = 0

LIMIT APIXNGluc >= 0

d/dt (ABIXNGluc) = - artP1 - artG1 - artF1 + venF1 - artR1 + venR1 - artS1 + venS1 - artK1 + venK1 + venL1 - artL1 - artU1 + venU1

INIT ABIXNGluc = 0

LIMIT ABIXNGluc >= 0

d/dt (AFIXNGluc) = + artF1 - venF1

INIT AFIXNGluc = 0

LIMIT AFIXNGluc >= 0

d/dt (ARIXNGluc) = + artR1 - venR1

INIT ARIXNGluc = 0

LIMIT ARIXNGluc >= 0

d/dt (ASIXNGluc) = + artS1 - venS1

INIT ASIXNGluc = 0

LIMIT ASIXNGluc >= 0

d/dt (AKIXNGluc) = + artK1 - venK1 - urineiXNGluc

INIT AKIXNGluc = 0

LIMIT AKIXNGluc >= 0

d/dt (IXNGlucExUr) = + urineiXNGluc

INIT IXNGlucExUr = 0

LIMIT IXNGlucExUr >= 0

d/dt (AUIXNGluc) = + artU1 - venU1

INIT AUIXNGluc = 0

LIMIT AUIXNGluc >= 0

{Flows}

FLIXNGluc = VmaxLIXNGlucs*CVLIXN/(KmLIXNGluc+CVLIXN)

FGIXNGluc = VmaxGIXNGlucs*CVGIXN/(KmGIXNGluc+CVGIXN)

artP1 = QP*CBIXNGluc

venP1 = QP*CVPIXNGluc

venG1 = QG*CVGIXNGluc

artG1 = QG*CBIXNGluc

artF1 = QF*CBIXNGluc

venF1 = QF*CVFIXNGluc

artR1 = QR*CBIXNGluc

venR1 = QR*CVRIXNGluc

artS1 = QS*CBIXNGluc

venS1 = QS*CVSIXNGluc

artK1 = QK*CBIXNGluc

venK1 = QK*CVKIXNGluc

venL1 = QL*CVLIXNGluc

artL1 = QLA*CBIXNGluc

urineiXNGluc = GFR*CVBIXNGluc*60/1000

artU1 = QU*CBIXNGluc

venU1 = QU*CVUIXNGluc

{Submodel "PN"}

{Reservoirs}

d/dt (AL8PN) = + FLiXN8PNc + venG2 + artL2 - venL2 + venP2 - FL8PN + transLItoL8PN - FL8PN

INIT AL8PN = 0

LIMIT AL8PN >= 0

d/dt (AG8PN) = - venG2 + artG2 - FG8PN - FG8PN

INIT AG8PN = 0

LIMIT AG8PN >= 0

d/dt (AB8PN) = - artL2 - artG2 + venL2 - artP2 - artR2 + venR2 - artS2 + venS2 - artK2 + venK2 - artF2 + venF2 - artU2 + venU2

INIT AB8PN = 0

LIMIT AB8PN >= 0

d/dt (AP8PN) = + artP2 - venP2

INIT AP8PN = 0

LIMIT AP8PN >= 0

d/dt (AR8PN) = + artR2 - venR2

INIT AR8PN = 0

LIMIT AR8PN >= 0

d/dt (AS8PN) = + artS2 - venS2

INIT AS8PN = 0

LIMIT AS8PN >= 0

d/dt (AK8PN) = + artK2 - venK2 - urine8PN

INIT AK8PN = 0

LIMIT AK8PN >= 0

d/dt (AF8PN) = + artF2 - venF2

INIT AF8PN = 0

LIMIT AF8PN >= 0

d/dt (Eliminate8PN) = + FL8PN + FG8PN

INIT Eliminate8PN = 0

LIMIT Eliminate8PN >= 0

d/dt (IXN8PNExUr) = + urine8PN

INIT IXN8PNExUr = 0

LIMIT IXN8PNExUr >= 0

d/dt (ALI8PN) = - transLItoL8PN + transSItoLI8PN

INIT ALI8PN = 0

LIMIT ALI8PN >= 0

d/dt (ASI8PN) = - transSItoLI8PN

INIT ASI8PN = AODOSE8PN

LIMIT ASI8PN >= 0

d/dt (AU8PN) = + artU2 - venU2

INIT AU8PN = 0

LIMIT AU8PN >= 0

{Flows}

FLiXN8PNc = VmaxLiXN8PNs*CVLIXN/(KmLiXN8PN+CVLIXN)

venG2 = QG*CVG8PN

artL2 = QLA*CB8PN

artG2 = QG*CB8PN

venL2 = QL*CVL8PN

artP2 = QP*CB8PN

venP2 = QP*CVP8PN

artR2 = QR*CB8PN

venR2 = QR*CVR8PN

artS2 = QS*CB8PN

venS2 = QS*CVS8PN

artK2 = QK*CB8PN

venK2 = QK*CVK8PN

artF2 = QF*CB8PN

venF2 = QF*CVF8PN

FL8PN = VmaxL8PNs* CVL8PN/(KmL8PN + CVL8PN)

FG8PN = VmaxG8PNs* CVG8PN/(KmG8PN + CVG8PN)

urine8PN = GFR*CVB8PN*60/1000

transLItoL8PN = Kb8PN*CLI8PN

transSItoLI8PN = Ka8PN*CSI8PN

artU2 = QU*CB8PN

venU2 = QU*CVU8PN

{Submodel "PNGluc"}

{Reservoirs}

d/dt (AP8PNGluc) = + artP3 - venP3

INIT AP8PNGluc = 0

LIMIT AP8PNGluc >= 0

d/dt (AL8PNGluc) = + venP3 + venG3 - venL3 + FL8PN

INIT AL8PNGluc = 0

LIMIT AL8PNGluc >= 0

d/dt (AG8PNGluc) = + artG3 - venG3 + FG8PN

INIT AG8PNGluc = 0

LIMIT AG8PNGluc >= 0

d/dt (AF8PNGluc) = + artF3 - venF3

INIT AF8PNGluc = 0

LIMIT AF8PNGluc >= 0

d/dt (AR8PNGluc) = + artR3 - venR3

INIT AR8PNGluc = 0

LIMIT AR8PNGluc >= 0

d/dt (AU8PNGluc) = + artU3 - venU3

INIT AU8PNGluc = 0

LIMIT AU8PNGluc >= 0

d/dt (AS8PNGluc) = + artS3 - venS3

INIT AS8PNGluc = 0

LIMIT AS8PNGluc >= 0

d/dt (AK8PNGluc) = + artK3 - venK3 - urine8PNGluc

INIT AK8PNGluc = 0

LIMIT AK8PNGluc >= 0

d/dt (AB8PNGluc) = - artP3 - artG3 - artF3 - artR3 - artS3 - artU3 - artK3 + venL3 + venF3 + venR3 + venS3 + venU3 + venK3

INIT AB8PNGluc = 0

LIMIT AB8PNGluc >= 0

d/dt (PNGlucEx) = + urine8PNGluc

INIT PNGlucEx = 0

LIMIT PNGlucEx >= 0

{Flows}

artP3 = QP*CB8PNGluc

artG3 = QG*CB8PNGluc

artF3 = QF*CB8PNGluc

artR3 = QR*CB8PNGluc

artS3 = QS*CB8PNGluc

artU3 = QS*CB8PNGluc

artK3 = QK*CB8PNGluc

venP3 = QP*CVP8PNGluc

venG3 = QG*CVG8PNGluc

venL3 = QL*CVL8PNGluc

venF3 = QF*CVF8PNGluc

venR3 = QR*CVR8PNGluc

venS3 = QS*CVS8PNGluc

venU3 = QU*CVU8PNGluc

venK3 = QK*CVK8PNGluc

urine8PNGluc = GFR*CVB8PNGluc*60/1000

FG8PN = VmaxG8PNs* CVG8PN/(KmG8PN + CVG8PN)

FL8PN = VmaxL8PNs* CVL8PN/(KmL8PN + CVL8PN)
